# The gatekeeper to gastric cancer; gastric microbiota invade the lamina propria in *Helicobacter pylori-*associated gastric carcinogenesis

**DOI:** 10.1101/2024.04.22.590522

**Authors:** Harriet J. Giddings, Ana Teodósio, Jack L. McMurray, Kelly Hunter, Zainab Abdawn, Jeffrey A. Cole, Claire D. Shannon-Lowe, Amanda E. Rossiter-Pearson

## Abstract

Stomach cancer is the fourth leading cause of cancer-related deaths worldwide. *Helicobacter pylori* is the main risk factor for gastric adenocarcinoma (GAC), yet the mechanism underpinning this association remains uncharacterised. Gastric intestinal metaplasia (GIM) represents the pre-cancerous stage and follows *H. pylori-*associated chronic gastritis (CG). Sequencing studies have revealed fewer *H. pylori* and more non-*H. pylori* bacteria in GAC. However, the spatial organisation of the gastric microbiota in health and disease is unknown. Here, we have combined RNA *in situ* hybridisation and immunohistochemistry to detect *H. pylori*, non-*H. pylori* bacteria and host cell markers (E-cadherin, Mucins 5AC and 2) from patients with CG (n=9), GIM (n=12), GAC and normal tissue adjacent to tumours (NATs) (n=3). Quantitative analysis of whole slide scans revealed significant correlations of *H. pylori* and other bacteria in CG and GIM samples. In contrast to sequencing studies, significantly fewer non-*H. pylori* bacteria were detected in *H. pylori-*negative patients. Importantly, whilst *H. pylori* exclusively colonised the gastric glands, non-*H. pylori* bacteria invaded the lamina propria in 3/4 CG and 5/6 GIM *H. pylori*-positive patients. Bacterial invasion was observed in 3/3 GAC samples and at higher levels than matched NATs. We propose that *H. pylori* ‘holds the keys’ to disrupt the gastric epithelial barrier, facilitating the opportunistic invasion of non-*H. pylori* bacteria to the lamina propria. Bacterial invasion could be a significant driver of inflammation in *H. pylori-*associated carcinogenesis. This proposed mechanism would both explain the synergistic roles of *H. pylori* and other bacteria and redirect attempts to prevent, diagnose and treat GAC.

## Introduction

Gastric adenocarcinoma (GAC) is the fourth leading cause of cancer-related deaths, accounting for 7.7% of cancer mortalities worldwide (1). *Helicobacter pylori* infection is associated with approximately 70% of GAC cases and the global prevalence of *H. pylori* infection is 50% (1). Correa’s cascade describes the histopathological changes to the gastric mucosa during progression to GAC (2). *H. pylori* first triggers chronic gastritis (CG), which can lead to gastric intestinal metaplasia (GIM), dysplasia and finally GAC. Although genotypes of *H. pylori* that encode the virulence factors CagA, VacA and HtrA (3) are more associated with severe disease outcomes, the mechanism by which a minority of *H. pylori* infections (1%) develop GAC remains unknown. In recent years, multiple 16S ribosomal RNA (16S rRNA) and metagenomic sequencing studies have profiled the human gastric microbiota during health and *H. pylori-* associated disease (4-7). The proposed model is that the healthy human stomach harbours a distinct microbial community structure and that upon *H. pylori* infection, this community structure shifts towards a dominance of *H. pylori* in CG. As carcinogenesis progresses, multiple other bacterial species displace *H. pylori* and dominate the GAC microbiota. However, sequencing studies are often prone to issues with contamination (8), do not offer spatial resolution and typically do not differentiate between transient and persistent gastric bacteria (9). A longstanding question is whether the gastric microbiota play a causative or correlative role in GAC. Here, we provide high resolution spatial images of *H. pylori* and non-*H. pylori* bacteria in patients with *H. pylori*-positive or negative CG, GIM, GAC and normal tissue adjacent to tumour (NAT). Importantly, we provide direct evidence of non-*H. pylori* bacteria invading the lamina propria in all stages of Correa’s cascade, suggesting that non-*H. pylori* bacteria might play a pivotal role in the early stages of *H. pylori-*associated carcinogenesis.

## Results and Discussion

### Localisation of *H. pylori* and the gastric microbiota in carcinogenesis

Archived formalin-fixed paraffin-embedded gastric corpus tissue were retrieved from the Human Biomaterials Research Centre (n=21). The associated clinical pathology reports confirmed samples were *H. pylori-*positive CG (n=4) or GIM (n=6) and *H. pylori-*negative CG (n=5) or GIM (n=6). An independent pathologist scored all samples for inflammation (supplementary Table 1). Gastric corpus tissue was also obtained from surgical resections of GAC and NATs (n=3). To detect *H. pylori* or non-*H. pylori* bacteria, sections were stained using RNAScope *in situ* hybridisation (ISH) probes against *H. pylori-*specific or conserved sequences of the 16S rRNA gene, respectively. The probe ‘Eubacteria’ was used to detect non-*H. pylori* bacteria. Immunohistochemistry enabled detection of the host cell markers E-cadherin, Mucins 5AC (Muc5AC) and 2 (Muc2) using automated tissue staining for CG and GIM samples. Manual tissue staining was used for GAC and NAT. Goblet cells, present only in GIM, secrete Muc2, whereas Muc5AC is found in both healthy and diseased gastric mucus. Representative images and quantification of indicated markers are shown in Figure 1A-D and Figure 1E-H, respectively. Consistent with previous studies (10, 11), *H. pylori* exclusively colonised the gastric glands (Figure 1B). The presence of Eubacteria correlated with *H. pylori* infection in CG and GIM, but were significantly reduced in the absence of *H. pylori* infection (n=11) (Figure 1F). This contrasts with the current dogma that there is a unique microbiota in *H. pylori-*negative individuals (4). Comparable levels of Muc5AC were observed across the patient groups and, as expected, there was a significant increase in Muc2 in GIM patients (Figure 1 G&H). Eubacteria increases in GAC compared with matched NAT samples, whilst *H. pylori* increases in NAT compared to GAC (Figure 1E&F). Although this is not statistically significant, this is in agreement with sequencing studies that show a displacement of *H. pylori* by gastric microbiota in GAC tissue (5, 7).

**Figure 1.**
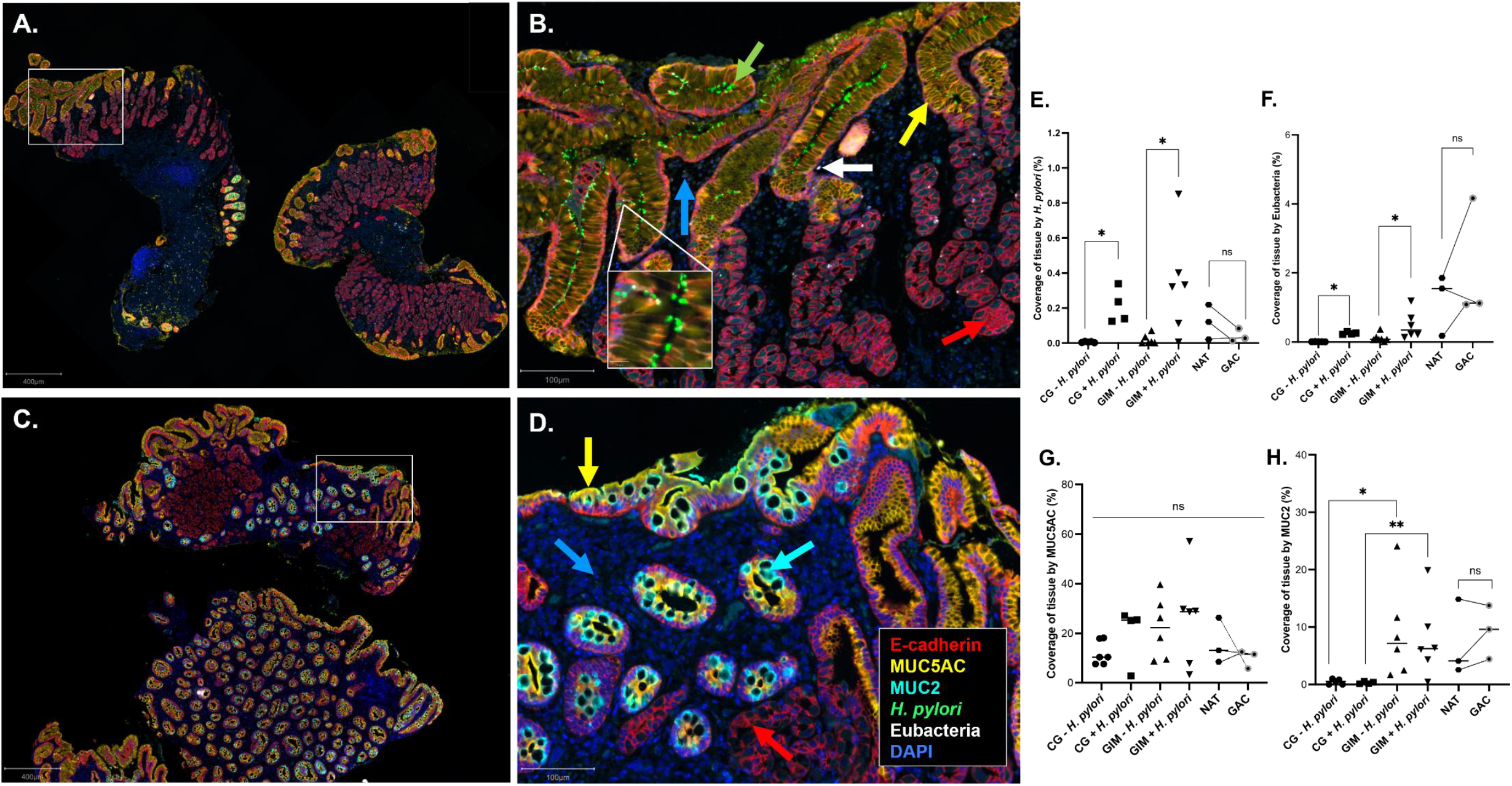
Quantification and spatial localisation of *H. pylori* and non-*H. pylori* bacteria in CG, GIM, NAT and GAC revealed by RNAScope *in situ* hybridisation (ISH) and immunohistochemistry (IHC). Whole slide scans of stained patient tissue sections were obtained using a Vectra whole slide scanner. Images were spectrally unmixed, viewed and quantified using QuPath. **A & C)** A representative whole slide scan of a GIM *H. pylori-*positive (A) or *H. pylori-*negative (C) section showing RNAScope ISH probes ‘H. pylori’ and ‘Eubacteria’ to detect *H. pylori* (green) and non-*H. pylori* bacteria (white), respectively. IHC staining against E-cadherin (red), Muc5AC (yellow) and Muc2 (turquoise) are also shown. **B & D)** A higher magnification of panel A and C, respectively. The coloured arrows show the indicated markers. **E-H)** Quantification of target markers as percentage coverage of tissue from whole slide scans for each patient group for *H. pylori* (E), Eubacteria (F) Muc5AC (G) and Muc2 (H). Lines between NAT and GAC data points show matched patient pairs. Horizontal bars indicate the median for each indicated patient group. A Mann-Whitney test was used to determine significance where * indicates *p*<0.05 and ** indicates *p*<0.01.

### *Helicobacter pylori* is associated with invasion of non-*H. pylori* bacteria to the lamina propria

Bacterial invasion is associated with multiple diseases of the gastrointestinal tract, causing damage to the epithelial architecture by triggering inflammatory responses (12, 13). Whilst *H. pylori* colonised the gastric glands, non-*H. pylori* bacteria invaded the lamina propria in 3/4 patients with *H. pylori-*associated CG and 5/6 *H. pylori-*positive patients with GIM (Figure 2A-C). In GAC, Eubacteria appear either evenly distributed across the tissue in two GAC patients (Figure 2D) or as dense regions of colonisation in one GAC patient (Figure 2E). To note, representative images of patients with *H. pylori-*negative CG or GIM (Figure 2 F&G, respectively) are shown. Very few Eubacteria are detected. It is most likely that these rare occurrences of Eubacteria in the stomach represent transient bacteria, rather than stable colonisation (9). Qualitative scoring of Eubacterial invasion showed a progressive increase of Eubacterial invasion from CG to GIM and GAC. Increased invasion was also observed in GAC compared with matched NATs (Figure 2H).

**Figure 2.**
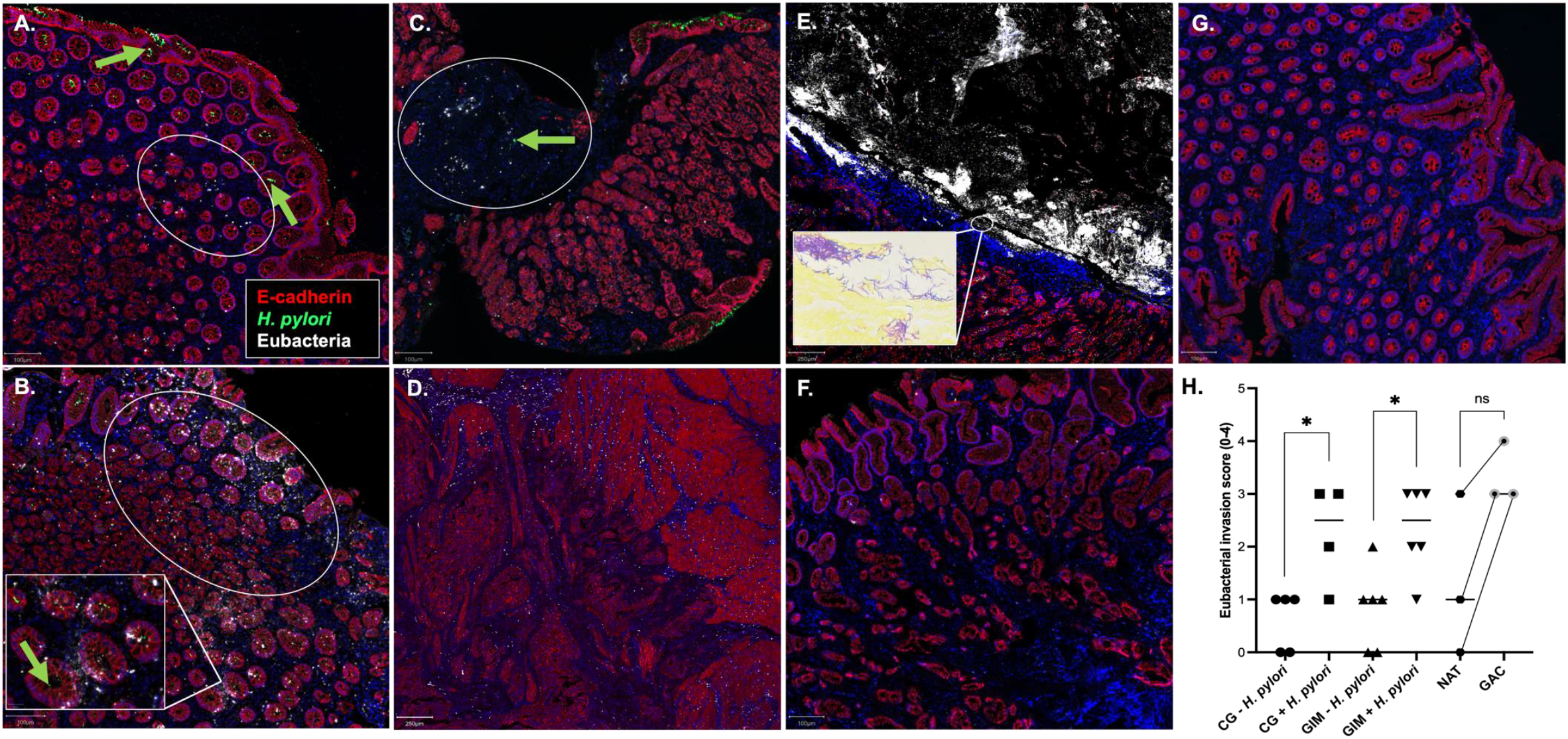
Non-*H. pylori* bacteria invade the gastric lamina propria in CG, GIM and GAC. Whole slide scans were obtained using a Vectra whole slide scanner. Images were spectrally unmixed and viewed using QuPath. **A-G)** Representative images showing Eubacterial (white) invasion in patients with CG (A-B), GIM (C), GAC (D&E), *H. pylori-*negative CG (F) and GIM (G) (Inset in Panel E shows bacterial cells in a modified Gram stain). E-cadherin (red), *H. pylori* (green), Eubacteria (white) and DAPI (blue) are shown for visualisation purposes. Arrows indicate the target markers. Eubacterial invasion is highlighted in white ovals for CG and GIM. A rare occurrence of *H. pylori* invasion to the lamina propria is indicated by the green arrow in (C). **H)** Qualitative FISH scoring of Eubacterial invasion for each patient group; 0 = no invasion, 1 = sparse, 2 = moderate, 3 = high and 4 = colonisation. Horizontal bars indicate the mean for each indicated patient group. A Mann-Whitney test was used to determine significance where * indicates *p*<0.05. Lines between NAT and GAC data points show matched patient pairs.

Next, bacterial invasion was confirmed at the cellular level using a modified Gram stain, which shows a mix of spindle shaped Gram-positive and Gram-negative bacteria in GAC (Figure 2E) and Gram-positive cocci in selected CG and GIM samples (data not shown). Further work is required to identify these invasive bacteria and the associated immune response. Invasion of *H. pylori* into the lamina propria is very rare across all patient samples, yet an instance of this can be seen in Figure 2C.

Recently, Sharafutdinov and colleagues reported that *H. pylori* strains encoding the trimeric form of HtrA, which proteolytically cleaves the cell junction proteins occludin, claudin-8 and E-cadherin, are associated with a higher GAC risk (3).

We propose that *H. pylori* acts as the ‘gatekeeper’ via HtrA-mediated cleavage of cell junction proteins of the gastric epithelial cell barrier (3), leading to infiltration of Eubacteria to the lamina propria (Figure 3). The prevalence of invasive bacteria in larger patient cohorts must be determined. Clinically, it should be trialled whether antibiotic eradication of *H. pylori* clears invasive bacteria, particularly in patients with GIM where eradication does not significantly reduce GAC risk. The role of invasive bacteria should be explored for translation into the prevention, diagnosis and treatment of GAC.

**Figure 3.**
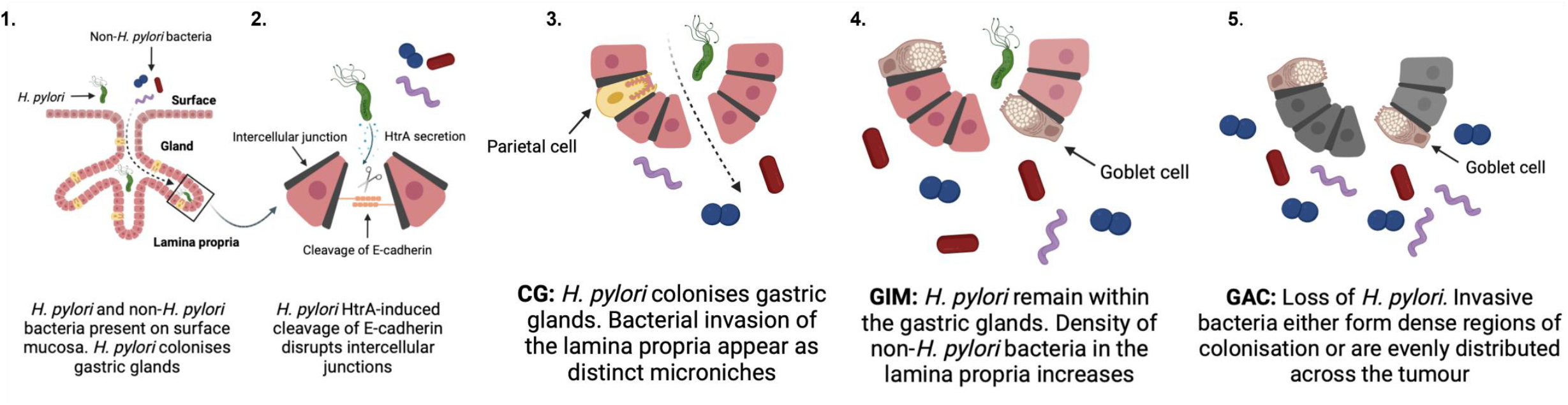
Illustrative model proposing *H. pylori* as the ‘gatekeeper’ for bacterial invasion to the gastric lamina propria. Stages 1-5 are (1) Eubacteria and *H. pylori* in the gastric surface and gland, respectively. (2) *H. pylori* secretes HtrA, which cleaves tight junction protein E-cadherin. Stages 3-5 are as described.

## Materials and Methods

Archived human gastric biopsies were sectioned and stained as described in the extended methods. Images were acquired using a Vectra Polaris whole slide scanner and analysed on open source QuPath software (14).

## Author Contributions

Conceptualisation: C.D.S-L. & A.E.R-P. Methodology: H.J.G, A.T., J.L.M, & A.E.R-P. Data analysis: H.J.G., A.T., J.L.M., K.H., Z.A. & A.E.R-P. Writing: H.J.G., J.A.C. & A.E.R-P. Funding acquisition: A.E.R-P.

## Acknowledgments

We thank the Royal Society and the University of Birmingham Centre Development fund from Cancer Research UK for funding (A.E.R.). We thank patients who have donated samples to HBRC. We also thank Michael Russell at HBRC for patient sample retrieval and Dr. Gary Reynolds for sample sectioning.

## Supplementary material

### Extended methods

#### 2.1 Sample preparation and pre-treatment protocol for automated tissue staining

Punch biopsy tissue samples were collected from consenting patients at the Queen Elizabeth Hospital, Birmingham, fixed in formalin and prepared at the Human Biomaterial Resource Centre (HBRC), University of Birmingham by embedding in paraffin (Ethics #17-285). 4 μM thick tissue sections were then prepared for RNAScope, immunohistochemistry and H&E staining, using a Leica BOND RX Fully Automated Research Stainer. Prior to staining, sections were prepared in the BOND RX, following three short protocols: 1) Deparaffinisation, rehydration, hydrogen peroxide and distilled water wash 2) Heating to 100°C in target retrieval buffer ER2 (pH 9; AR9640) for 45 min and manual washing with distilled water followed by 100% ethanol and drying at 60°C. 3) Incubation in protease III solution for 30 min, followed by a final wash in distilled water.

#### 2.2 Optimisation of RNAScope treatments and immunohistochemistry antibody concentrations for 5-plex automated staining

Automated RNAScope C1 and C2 probes against *H. pylori* (ACD-Bio, #542938) and Eubacteria (ACD-Bio; #464468), respectively, were first tested on healthy colon tissue in a fluorescent multiplex assay using an RNAScope LS Multiplex Fluorescent Reagent Kit (ACD Bio; 322800), following the standard protocol recommended for this platform. Pre-treatment of tissue slides for automated tissue staining with lysozyme was not used as this was found to affect tissue integrity. However, automated tissue staining with lysozyme (micro bacteria detection protocol provided by ACD-Bio) and a 3-plex panel (E-Cadherin and RNAScope probes against Eubacteria and *H. pylori*) was used on all samples (n=21) to ensure bacterial signal was not underestimated in comparison to 5-plex stained images (data not shown).

A positive and negative control probe section was used on every automated RNAScope run to validate and assess its quality and the sensitivity of the assay. The bacterial gene DapB (ACD Bio; 312038) was used as negative control to confirm the absence of background noise and a cocktail of housekeeping genes Polr2A C1, PPIB C2 and UBC C3 (ACD Bio; 320868) were used as positive control to validate the detection of the signal and the tissue integrity. *H. pylori* gene sequences can bind to both 16S rRNA probes against *H. pylori* and Eubacteria, whilst non-*H. pylori* bacteria stain with only the Eubacteria probe.

All antibodies used in IHC steps were optimised prior, using chromogenic DAB staining of healthy colon tissue in addition to a Leica Bond Polymer Refine Detection kit (Leica; DS9800). Antigen retrieval was tested using pH6 (Leica Bond TM Epitope Retrieval 1; AR9961) and pH9 (Leica Bond TM Epitope Retrieval 2; AR9640) buffers by heating to 100°C for 20 min. Three different dilutions were tested for each antibody, as recommended by manufacturer. Ideal staining pattern and intensity was assessed and approved by a pathologist, whereby slides were then used as reference throughout the validation process. All antibodies were then tested for compliance with the RNAScope pre-treatment, to ensure stability after protease III digestion. Once each antibody was assigned OPAL fluorophore single fluorescence assays were directly compared against DAB-stained colon tissue to optimise OPAL concentration. To assess epitope stability following heat steps and to define the order of the antibodies in the multiplex sequence, each antibody was tested individually in the different positions of the panel. Further probe and antibody information can be found in Table 1.

**Table 1.** Automated 5-plex RNAScope probe and IHC antibody information.

| Marker | Marker type | Supplier | Clone | Cat. No. | Marker dilution | Fluorophore | Fluorophore cat. No. | Fluorophore dilution |
| --- | --- | --- | --- | --- | --- | --- | --- | --- |
| *H. pylori* | RNAscope C1 | ACD | N/A | N/A | RTU | 520 | FP1487001KT | 1:150 |
| Eubacteria | RNAscope C2 | ACD | N/A | N/A | 1:50 | 620 | FP1495001KT | 1:150 |
| E-cadherin | Antibody | Cell Signalling Technology | 24E10 | 3195S | 1:250 | 690 | FP1497001KT | 1:200 |
| MUC2 | Antibody | Novus Biologicals | Polyclonal | NBP1-31231 | 1:1000 | 480 | FP1500001KT | 1:150 |
| MUC5AC | Antibody | Invitrogen | 2-25LE | MA119346 | 1:1000 | 570 | FP1488001KT | 1:150 |

#### 2.3 Automated 5-plex co-RNAScope *in situ* hybridisation/immunohistochemistry protocol

The automated RNAScope Multiplex Fluorescent LS assay (ACD Bio, 322800) was conducted according to manufacturer’s instructions, using C1 and C2 probes against *H. pylori* and Eubacteria, respectively. An automated staining protocol was immediately followed using a Leica BOND RX, to fluorescently label E-cadherin, MUC5AC and MUC2. Images were acquired using a Vectra Polaris whole slide scanner. Exposure times on the Vectra Polaris Slide scanner for the DAPI, 480, 520, 570, 620 and 690 channels were 1.13 ms, 2.47 ms, 36.06 ms, 5.61 ms, 29.58 ms, and 7.46 ms, respectively.

#### 2.4 Manual 3-plex sample preparation and pre-treatment protocol

Gastrectomy tissue samples were collected from consenting patients at the Queen Elizabeth Hospital, Birmingham, fixed in formalin and prepared at the Human Biomaterial Resource Centre (HBRC), University of Birmingham by embedding in paraffin (IRAS ethics #276525 and 20/NW/0001). 4 μM thick tissue sections were then prepared for staining at the Institute of Cancer and Genomic Sciences, University of Birmingham. Prior to staining, sections were prepared according to the manual RNAscope v2 assay (ACD Bio; 323110). Antigen retrieval was conducted in in Simport Easy Dip staining jars (Simport; M900-12W) in a microwave oven for 1 minute at 100% power, followed by 15 minutes at 20% power with intervals to prevent boiling.

#### 2.5 Optimisation of RNAScope treatments and IHC antibody concentrations for manual 3-plex staining

Manual C1 (ACD Bio; 542931), C2 (ACD Bio; 464461-C2) and C3 (ACD Bio; 1112731-C3) probes were assigned to *H*. pylori and Eubacteria, respectively and prepared according to manufacturer’s instructions. Probes were tested alone and in combination with the micro bacteria detection protocol provided by ACD, including addition of lysozyme, using *in vitro* infection models. All antibodies used in IHC steps were optimised prior by the automated staining protocol and tested manually *in vitro* by staining healthy human gastric organoid models. Antigen retrieval pH and method was optimised in a water bath or microwave oven at ∼98°C for 20-45 min to heat sections. pH6 (Perkin Elmer) and pH9 (Perkin Elmer) buffers were also compared during each heating method.

#### 2.6 Manual co-RNAScope ISH/IHC protocol

The RNAScope multiplex fluorescent v2 assay (ACD Bio; 323110) was conducted according to manufacturer’s instructions; all incubation periods were at 40°C in a humidified incubator. RNAScope multiplex fluorescent v2 assay, C1 and C2 probes were used for *H. pylori* and Eubacteria. Sections were immediately washed and blocked for 45 min in blocking buffer containing 0.3% Triton-X, 3% BSA, 0.3% Goat Serum and 1X PBS. Sections were washed, and E-cadherin primary antibody was added to sections over night at 4°C. After 16-20 h, sections were washed and then incubated with secondary antibody Alexa Fluor 594, diluted 1:1000 in blocking buffer, for 1 h at room temperature. Exposure times varied between samples and were determined at the time of scanning.

#### 2.7 Manual 3-plex IHC protocol

Manual IHC staining of E-cadherin, MUC5AC and MUC2 was conducted on prepared sections using the Opal 7-colour manual IHC Kit (Perkin Elmer; NEL811001T), according to manufacturer’s instructions. Exposure times varied between samples and were determined at the time of scanning. Antibody information and corresponding fluorophores are detailed in Table 4.

**Table 3.** Manual RNAScope probe and IHC antibody information.

| Marker | Marker type | Supplier | Clone | Cat. No. | Marker | Fluorophore | Fluorophore cat. | Fluorophore |
| --- | --- | --- | --- | --- | --- | --- | --- | --- |

|  |  |  |  |  | dilution |  | no. | dilution |
| --- | --- | --- | --- | --- | --- | --- | --- | --- |
| *H. pylori* | RNAScope C1 | ACD | N/A | 542931 | RTU | TSA vivid 520 | 7526 | 1:1500 |
| Eubacteria | RNAScope C2 | ACD | N/A | 464461-C2 | 1:50 | TSA vivid 570 | 7523 | 1:1500 |
| E-cadherin | Antibody | Cell Signalling Technology | 24E10 | 3195S | 1:400 | Alexa Fluor 594 | A20185 | 1:1000 |

**Table 4.**
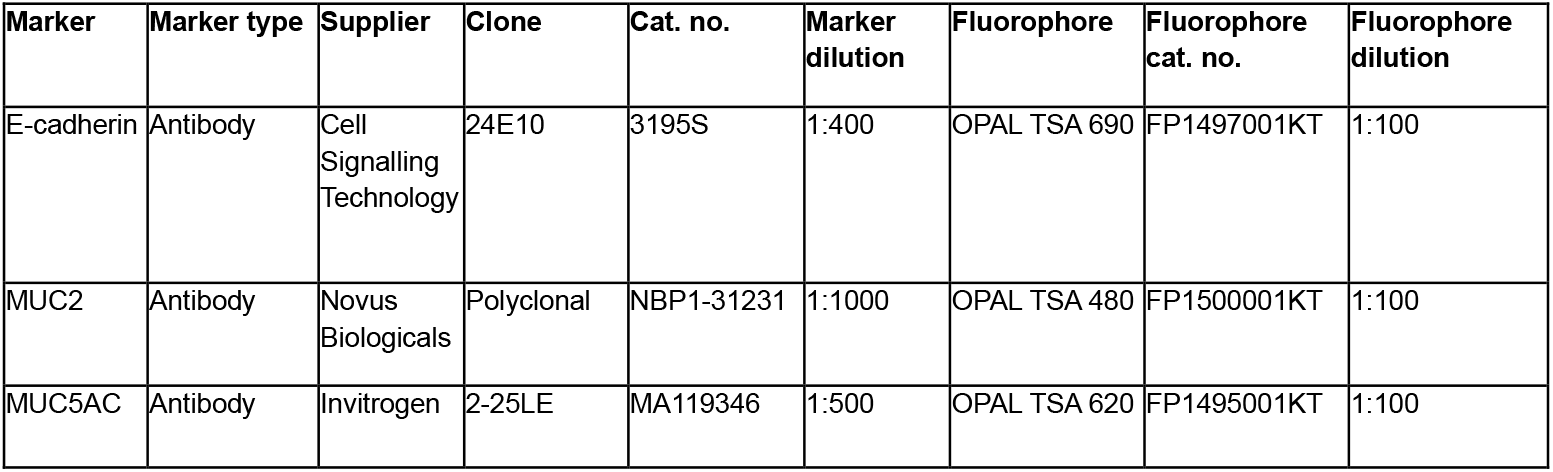
Manual 3-plex IHC antibody information.

#### 2.8 Counterstaining and section visualisation

All sections were counterstained with spectral DAPI (Akoya Biosciences) mounted with ProLong Diamond Antifade Mountant according to manufacturer’s instructions. All mounted sections were stored in the dark at 4°C until viewing. During optimization steps, sections were visualised using a Zeiss LSM 900 confocal microscope or Mantra 2 Quantitative Pathology Digital Workstation (Akoya Biosciences) and all final images were obtained on a Vectra Polaris Multispectral Whole Slide Scanner (Akoya Biosciences) and saved as a .qptiff file. Images were viewed at 40x magnification using Phenochart Whole Slide Viewer with PhenoImager HT (Akoya Biosciences) or QuPath software. .qptiff image files were spectrally unmixed by importing and stamping in Phenochart Whole Slide Viewer (Akoya Biosciences), unmixed and exported in InForm software (Akoya Biosciences) and finally restitched as a BIGTIFF file using Visiopharm software (Visiopharm, Hørsholm, Denmark).

#### 2.9 Qualitative image analysis

All images were viewed using Phenochart Whole Slide Viewer (Akoya Biosciences) or open source QuPath software (Version 0.4.3). The presence of each marker of interest (*H. pylori*, Eubacteria, E-Cadherin, MUC5AC and MUC2) were scored independently (H.J.G. and A.E.R-P.) whereby 1 = no/low staining intensity, 2 = medium staining intensity, 3 = high staining intensity. Invasion of Eubacteria was also qualitatively scored, whereby 0 = no invasion, 1 = sparse invasion, 2 = moderate invasion (patches of bacteria across sample), 3 = high invasion (multiple clear regions of bacterial invasion across sample) and 4 = dense colonisation of bacteria (large regions of sample colonised).

#### 2.10 Quantitative image analysis

Manual image analysis was conducted using the free and open source Qupath software (Version 0.4.3). For each whole-slide scan, tissue regions were annotated and defined as a region of interest (ROI). Tissue detection was then performed based on the average values of all channels using three separately created pixel thresholders. Each pixel thresholder was used to calculate tissue area (μM^2^), which was exported, and mean tissue area value was obtained. Three separate pixel thresholders were created for each individual OPAL channel, corresponding to - *H. pylori*, Eubacteria, MUC5AC or MUC2. The thresholders were saved, and average area annotation measurements (μM^2^) were obtained for each channel. To account for *H. pylori* double-staining and detection in both the *H. pylori* and Eubacteria channels, absolute *H. pylori* area was calculated by calculating total Eubacteria channel area minus combined *H. pylori* and Eubacteria channel areas. For each whole slide scan, average percentage area coverage of each marker of interest was calculated by dividing individual channel area over total tissue area x 100. Data were exported from Excel to GraphPad Prism 9 (Version 9.5.1). For statistical analysis, a Mann-Whitney test was performed, whereby * indicates *p*<0.05, ** indicates *p*<0.01 and ns indicates non-significant. Data are presented as median ± SEM, for each group.

#### 2.11 Additional staining

Additional sections were obtained and stained with H&E for further inflammation analysis and scoring by an independent pathologist. A modified Gram stain was conducted for staining of invasive bacteria in tissue sections of interest (15).

**Supplementary Table 1.** Patient demographics, clinical inflammatory scores and raw values of target markers as % coverage of whole slide scan.

| Patient | Patient group | Gender | Diagnosis | PPI use | H&E Inflammation grade | Invasion Score (0-4) | *H. pylori* coverage (%) | Eubacteria coverage (%) | MUC5AC coverage (%) | MUC2 coverage (%) |
| --- | --- | --- | --- | --- | --- | --- | --- | --- | --- | --- |
| 1 | CG *H. pylori* positive | Male | Mild active *H. pylori* associated gastritis | None | Moderate | 3 | 0.236 | 0.218 | 2.793 | 0.377 |
| 2 | CG *H. pylori* positive | Male | Active chronic gastritis | None | Mild | 1 | 0.139 | 0.234 | 25.531 | 0.107 |
| 3 | CG *H. pylori* positive | Male | Mild active *H. pylori* associated gastritis | None | Moderate | 3 | 0.339 | 0.303 | 25.238 | 0.123 |
| 4 | CG *H. pylori* positive | Male | Mild active chronic gastritis | Lansoprazole | Moderate | 2 | 0.125 | 0.256 | 27.231 | 0.578 |
| 5 | CG *H. pylori* negative | Male | Mild reactive gastropathy | None | Mild | 1 | 0.003 | 0.003 | 17.926 | 0.834 |
| 6 | CG *H. pylori* negative | Male | Mild chronic gastritis | None | Mild | 0 | 0.006 | 0.005 | 10.44 | 0.0176 |
| 7 | CG *H. pylori* negative | Male | Mild non-specific chronic inflammation | None | Mild | 1 | 0.008 | 0.001 | 10.51 | 0.018 |
| 8 | CG *H. pylori* negative | Male | Mild chronic inactive gastritis | Lansoprazole | Moderate | 1 | 0.009 | 0.004 | 18.167 | 0.992 |
| 9 | CG *H. pylori* negative | Male | Corrobative of inflamed Barretts mucosa | Lansoprazole | Moderate | 0 | 0.004 | 0.007 | 7.662 | 0.534 |
| 10 | GIM *H. pylori* positive | Male | Severe active chronic infection with gastritis/atrophy | None | Severe | 2 | 0.112 | 0.128 | 7.856 | 10.028 |
| 11 | GIM *H. pylori* positive | Male | Chronic active gastritis and intestinal metaplasia | None | Severe | 3 | 0.005 | 0.217 | 57.06 | 4.321 |
| 12 | GIM *H. pylori* positive | Male | Mild active *H. pylori* associated gastritis and intestinal metaplasia | None | Patchy severe | 3 | 0.401 | 0.472 | 3.266 | 19.861 |
| 13 | GIM *H. pylori* positive | Male | *H. pylori* associated gastritis and intestinal metaplasia | None | Moderate | 1 | 0.85 | 0.003 | 28.602 | 6.103 |
| 14 | GIM *H. pylori* positive | Male | Intestinal metaplasia with squamous and columnar mucosa | Lansoprazole | Moderate | 2 | 0.332 | 0.214 | 29.725 | 0.419 |
| 15 | GIM *H. pylori* positive | Male | *H. pylori* associated gastritis and intestinal metaplasia | None | Mild | 3 | 0.321 | 0.685 | 29.032 | 6.36 |
| 16 | GIM *H. pylori* negative | Male | Chronic gastritis and intestinal metaplasia | None | Moderate | 0 | 0.003 | 0.075 | 39.776 | 11.663 |
| 17 | GIM *H. pylori* negative | Male | N/C | None | Mild | 0 | 0.004 | 0.082 | 8.84 | 6.16 |
| 18 | GIM *H. pylori* negative | Female | Mild active gastritis and focal intestinal metaplasia | None | *Missing | 1 | 0.008 | 0.01 | 9.479 | 24.095 |
| 19 | GIM *H. pylori* negative | Female | Active chronic gastritis and intestinal metaplasia | None | *Missing | 1 | 0.036 | 0.07 | 31.402 | 2.499 |
| 21 | GIM *H. pylori* negative | Male | Focal active chronic gastritis and intestinal metaplasia/atrophy | None | Moderate | 1 | 0.003 | 0.073 | 18.264 | 1.698 |
| 22 | GIM *H. pylori* negative | Male | Mild chronic gastritis and intestinal metaplasia | Rabeprazole | Mild | 2 | 0.072 | 0.138 | 26.334 | 8.198 |
| 23 | GAC | Male | Gastric adenocarcinoma | N/A | N/A | 4 | 0.22 | 4.163 | 12.525 | 9.595 |
| 23 | NAT | Male | Healthy tissue adjacent to tumour | N/A | N/A | 3 | 0.084 | 1.853 | 8.49 | 4.084 |
| 24 | GAC | Male | Gastric adenocarcinoma | N/A | N/A | 3 | 0.015 | 1.192 | 5.839 | 13.749 |
| 24 | NAT | Male | Healthy tissue adjacent to tumour | N/A | N/A | 0 | 0.028 | 0.175 | 26.362 | 14.853 |
| 25 | GAC | Male | Gastric adenocarcinoma | N/A | N/A | 3 | 0.021 | 1.122 | 11.587 | 4.441 |
| 25 | NAT | Male | Healthy tissue adjacent to tumour | N/A | NA | 1 | 0.114 | 1.58 | 13.18 | 2.553 |

